# A tale of two plasmids: contributions of plasmid associated phenotypes to epidemiological success among *Shigella*

**DOI:** 10.1101/2021.12.17.473221

**Authors:** P. Malaka De Silva, George E. Stenhouse, Grace A. Blackwell, Rebecca J. Bengtsson, Claire Jenkins, James P.J. Hall, Kate S. Baker

## Abstract

Dissemination of antimicrobial resistance (AMR) genes by horizontal gene transfer (HGT) mediated through plasmids is a major global concern. Genomic epidemiology studies have shown varying success of different AMR plasmids during outbreaks, but the underlying reasons for these differences are unclear. Here, we investigated two *Shigella* plasmids (pKSR100 and pAPR100) that circulated in the same transmission network but had starkly contrasting epidemiological outcomes to identify plasmid features that may have contributed to the differences. We used plasmid comparative genomics to reveal divergence between the two plasmids in genes encoding AMR, SOS response alleviation, and conjugation. Experimental analyses revealed that these genomic differences corresponded with reduced conjugation rates for the epidemiologically successful pKSR100, but more extensive AMR, reduced fitness costs, and a reduced SOS response in the presence of antimicrobials, compared with the less successful pAPR100. The discrepant phenotypes between the two plasmids are consistent with the hypothesis that plasmid associated phenotypes contribute to determining the epidemiological outcome of AMR HGT and suggest that phenotypes relevant in responding to antimicrobial pressure and fitness impact may be more important than those around conjugation in this setting. Plasmid phenotypes could thus be valuable tools in conjunction with genomic epidemiology for predicting AMR dissemination.

## Introduction

Antimicrobial resistance (AMR) is a pressing global public health crisis. Bacterial pathogens become resistant to antimicrobials through either chromosomal mutations or by acquiring new AMR determinants through horizontal gene transfer (HGT) of mobile genetic elements (MGEs), such as plasmids (1, 2). HGT plays an important part in the dissemination of AMR genes, evidenced by multiple reports of AMR emergence of similar AMR genes from different locations around the world. For example, plasmid-mediated colistin resistance conferred by the *mcr-1* gene was first identified in China in 2011 and has since spread across five continents (3) and many Gram-negative bacterial species by virtue of its being carried on a plasmid capable of inhabiting multiple hosts (4). Similarly, *Klebsiella pneumoniae* carbapenemases (KPCs), which were originally observed in the USA in 1996 (5, 6), and CTX-M extended spectrum beta lactamases (ESBL), which were thought to be mobilised from the chromosome of *Kluyvera* spp., have been reported in multiple geographical regions as a result of AMR plasmid dissemination (7-10). Owing to the rapid and extensive dissemination of AMR genes through HGT, it is critical to understand how different plasmid ‘vehicles’ affect the behaviour and spread of AMR genes.

Headway has been made towards understanding AMR epidemiology with reference to the specific plasmids that carry AMR determinants for a number of pathogen-plasmid combinations, largely from hospital settings (11-15). *in vitro* work as well as modelling studies — in some cases supported by epidemiological data — suggest that the phenotypes of plasmids, such as plasmid fitness costs, resistance gene profile, and other plasmid-conferred traits, may have a role in driving transmission and persistence of AMR across bacterial host populations (16-21). Recent studies have attempted to associate non-AMR phenotypes such as conjugation with pathogen and plasmid epidemiology, including in clinical settings (11, 12, 22). However, the drivers of AMR-HGT in non-hospital associated pathogens are likely to be distinct owing to the environmental differences (e.g. in antimicrobial pressures and transmission routes) and the comparatively under-observed bacterial populations.

Tracking plasmid-mediated AMR emergence in the community therefore requires a well-surveyed and characterised pathogen population. The Gram-negative diarrhoeal pathogen *Shigella* provides a highly observable community infection because infection is almost always symptomatic, the disease is reportable, and there is no substantial animal or environmental reservoir. Using genomic epidemiology, we recently investigated the emergence of AMR in the relatively closed transmission network of *Shigella* infections amongst men who have sex with men (MSM) in the United Kingdom (23, 24). The selective pressure caused by antimicrobial treatment of sexually transmitted infections (STIs) such as syphilis and gonorrhoea (common co-infections among shigellosis affected MSM (23, 25)), led to the global emergence of a sublineage of *S. flexneri* 3a following the acquisition plasmid pKSR100, encoding azithromycin resistance (23). This sustained selection pressure subsequently led to the convergent acquisition of multiple azithromycin resistance plasmids in multiple other *Shigella* sublineages, directly triggering and intensifying epidemic waves of Shigellosis (24, 26). The acquisition of different azithromycin-resistance plasmids by these different sublineages provides us with an epidemiologically grounded model to test why some plasmids proliferate in a population, and why some do not.

The two IncFII plasmids under study here derive from a cross-sectional analysis of UK *Shigella* strains detected during routine surveillance between 2008 and 2014. During this period, we demonstrated that *S. flexneri* 2a emerged within the MSM community, as two azithromycin resistant sublineages with markedly different epidemiology (24). The transient or ‘minor’ sublineage was estimated to have emerged earlier (most recent common ancestor [MRCA] in 1996) but only caused 7 cases during the study period, whereas the comparatively recent (MRCA 2011) persistent or ‘major’ sublineage, caused 49 cases in the study period, and disseminated internationally (24, 26, 27). Although both sublineages were azithromycin resistant, the plasmids conferring azithromycin resistance varied. The successful ‘major’ sublineage carried pKSR100 (24), a plasmid which has continued to spread globally throughout shigellae, while the other ‘minor’ sublineage carried azithromycin resistance on a different plasmid (herein termed pAPR100) which has failed to mirror the epidemiological success of pKSR100. Furthermore, pKSR100 has continued to acquire novel AMR genes including a multidrug resistance (MDR) integron (24) and, more recently, ESBL genes (28). As both pKSR100 and pAPR100 plasmids conferred the critical azithromycin resistance phenotype, we hypothesised that non-azithromycin resistance related plasmid phenotypes of these co-circulating plasmids may have contributed to their disparate epidemiological outcomes.

Thus here, we use these two plasmids to examine which plasmid phenotypes associate with the emergence of AMR in a community transmitting obligate pathogen. We extend the genomic epidemiological evidence for the respective global distribution of the plasmids and use a comparative genomics-guided approach to determine the discrepant phenotypes between these two co-circulating plasmids with markedly different epidemiological outcomes.

## Results

### pKSR100 has greater epidemiological success than pAPR100

As pKSR100 and pAPR100 had markedly different epidemiological success in the original study (conducted in the UK and France from 2008 – 2016) (24), we investigated whether the broader scientific research base and subsequent public health surveillance activity supported this discrepancy. In the original study, the pKSR100 carrying sublineage of *S. flexneri* 2a rapidly emerged as a dominant sublineage in a short space of time compared to the pAPR100 carrying sublineage even though the latter was circulating in the population prior to the acquisition of pKSR100 (24).

Since the original reporting of pKSR100 in Australia, Canada, France and the UK, further pKSR100 and pKSR100-like plasmids have been reported and/or deposited in the National Center for Biotechnology Information (NCBI) non-redundant database. Specifically, a BLASTn search returned 15 hits with over 95% query coverage and 95% identity from ≥ 7 *Shigella* serotypes across four continents, continuing to drive national and global shigellosis dynamics (26, 27, 29-32) (Figure 1A). Contrasting with the widespread dissemination of pKSR100, no BLAST hits with more than 87% query coverage for pAPR100 were returned, suggesting no distribution beyond its original detection in *S. flexneri* 2a in the UK. To extend the investigation of plasmid dissemination to unassembled genomes (such as those from routine public health surveillance activity), we screened both plasmids against the 661K COBS data structure, which provides a snapshot all bacterial genome data in the ENA as of November 2018 (33). This search detected kmer similarity matches of over 0.80 of pKSR100 in 1,926 publicly available bacterial genomes compared with only 46 bacterial genomes for pAPR100, further confirming the extensive dissemination of pKSR100. The genomes containing similarity matches for pKSR100 belonged to diverse bacterial hosts comprising 9 sequence types of *Shigella*, and 34 sequence types of *E. coli* and 2 other species (a *Klebsiella pneumoniae* ST258 and a *Salmonella enterica* ST11), while pAPR100 hits only came from 5 sequence types of *Shigella* and 13 sequence types of *E. coli* (Figure 1B). Having established that pKSR100 had ongoing epidemiological success compared with pAPR100 through these analyses, we then set out to compare phenotypes of the two plasmids.

**Figure 1.**
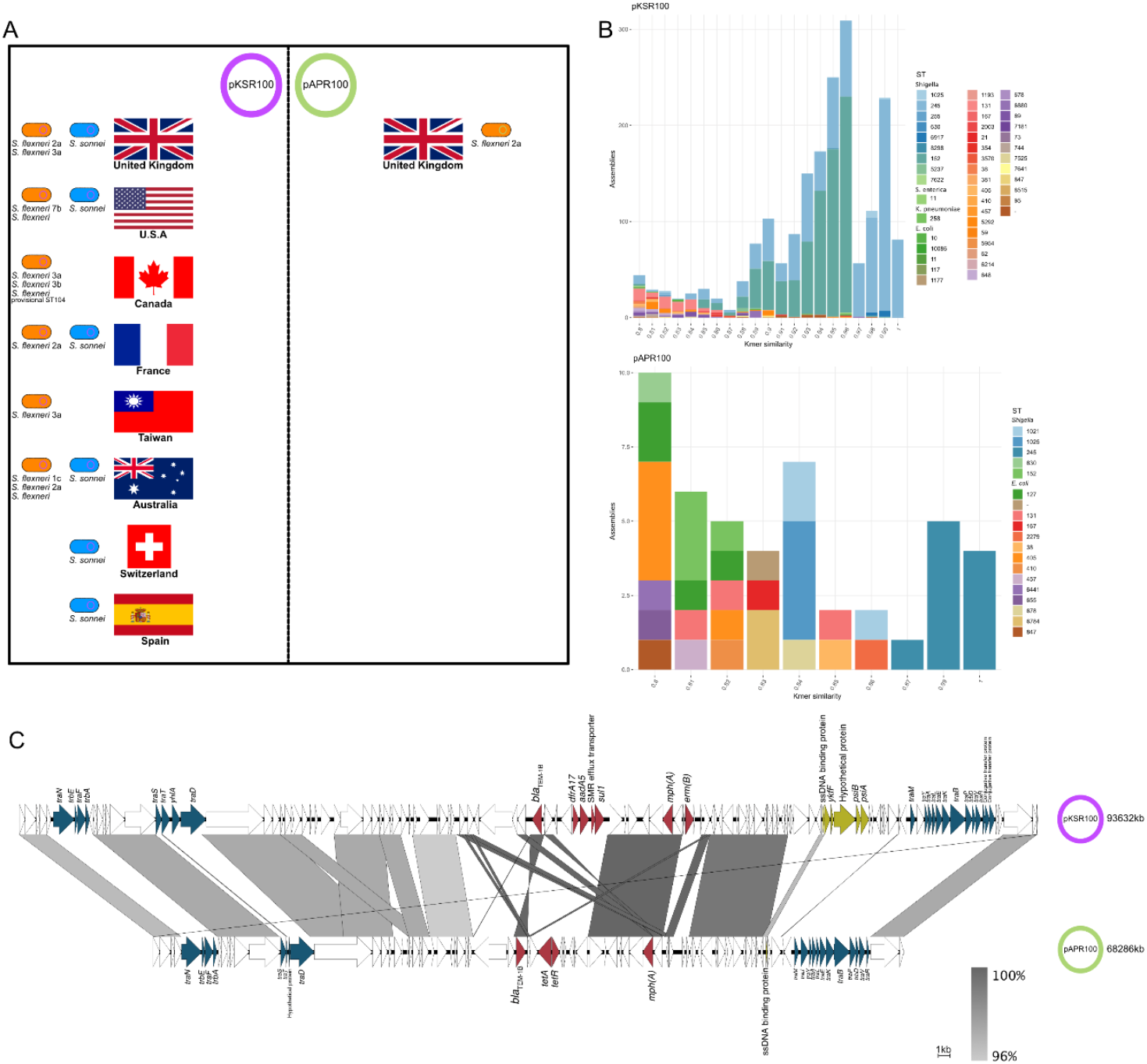
Relative global distribution and comparative genomics of pKSR100 & pAPR100. The disparate global and species distribution of pKSR100 and pAPR100 is shown as determined by (A) NCBI database BLAST and literature review where pKSR100 and pKSR100-like plasmids have been detected in and reported from eight countries and in multiple *Shigella* subtypes. In contrast, pAPR100 has only been detected in *S. flexneri* 2a in the UK in our previous study; (B) Species and sequence type distribution of genomes containing pKSR100 or pAPR100 in the 661K COBS data structure with kmer similarities of >0.80. (C) Comparison of the genetic content of both plasmids. Areas of synteny (using a cut off of 95% BLAST identity) are shown intervening grey bars coloured according to the inlaid legend. Regions with variation between the two plasmids are broadly categorised into three main groups; (i) conjugation machinery related (blue), (ii) SOS response alleviation related (green), and (iii) AMR related (red).

### pKSR100 and pAPR100 broadly differ across three gene categories

To guide our phenotypic experiments, we first performed comparative genomic analysis of the plasmid sequences to identify areas of differing genetic content. The plasmids had a high degree of conservation between their genetic content (Figure 1C, Supplementary file S2) with a total of 49 genes shared between both plasmids and 122 genes unique to each. However, the variances we observed with the gene content on both plasmids could be categorised into three broad functional categories; namely (i) conjugation machinery related genes, implying different conjugative abilities between pKSR100 and pAPR100 (ii) genes related to SOS response alleviation, indicating that perhaps pKSR100 conferred a fitness advantage related to stress responses and (iii) AMR genes, suggesting that perhaps the advantage of pKSR100 lay in conferring greater or broader resistance to antimicrobials (Figure 1C). Thus, we then proceeded to measure and compare the phenotypic differences between these two plasmids with disparate epidemiological outcomes. The two sequences of the pKSR100 and pAPR100 variants used in this study have been uploaded to the NCBI under accession numbers (TBP) and (TBP) respectively.

### pKSR100 is less conjugative than pAPR100 from its native donor

We observed variation in the conjugation related *tra* genes and *trb* genes between the two plasmids even though none of the genes were absent in either of the plasmids (Figure 1C). These variations at the nucleotide level were observed at a 95% BLAST identity cut-off between the two plasmid sequences. Given these discrepancies in conjugation related genes and the differential dissemination of the plasmids at an epidemiological level, we investigated the conjugation rates of pKSR100 and pAPR100 in shigellae and in model strain *E. coli* MG1655.

Both pKSR100 and pAPR100 carried T4SS genes required for conjugation and were conjugative from *S. flexneri* to, and between, *E. coli* MG1655 and a clinical isolate of *S. sonnei* (Figure 2). The *S. sonnei* clinical isolate we used, *S. sonnei* 216, originated from the same collection of isolates as the native hosts of pKSR100 and pAPR100 (24) but did not naturally carry either plasmid, and thus acts as a natural and relevant recipient. We measured conjugation efficiency (CE) across a number of variables including: from various donors (native and isogenic); to various recipients (*E. coli* and *S. sonnei* 216); in/on liquid and solid media; and across several time points (Figure 2, Figure S2).

**Figure 2.**
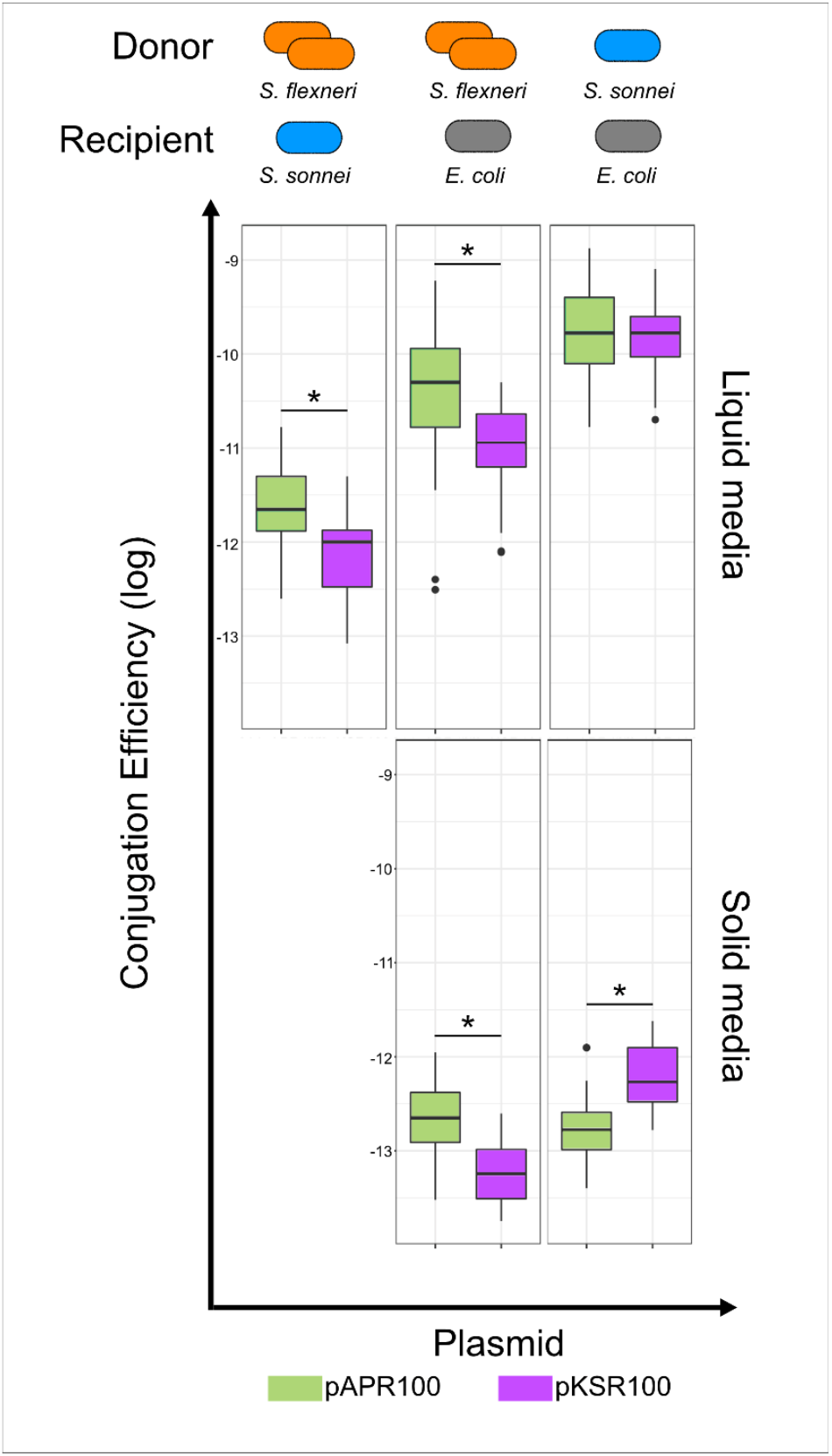
Conjugation efficiencies of pKSR100 and pAPR100 among different donors, recipients, and media conditions. Different donor strains included native *S. flexneri* hosts (orange) and clinical isolate *S. sonnei* 216 as an isogenic donor (blue) while the recipient strains were either *E. coli* MG1655 (in grey) or *S. sonnei* 216 (in blue). Results for each plasmid are coloured according to the inlaid key. Results from liquid media are shown upper while solid is shown lower. Each box plot represents the combined results of all the time points used in LM models, where there are 12 replicates (four biological, three technical) for each time point. The asterisks denote significance as determined LMs. The p-values for each of the panels from left to right are; top row – p=0.043, p<0.000, bottom row – p=0.026, p=0.025.

Generally, pAPR100 had a better conjugation efficiency compared to pKSR100 from their native donors (*i*.*e*. the major and minor sublineages of *S. flexneri* 2a (25)) into either *S. sonnei* 216 or *E. coli* MG1655, regardless of media type (post-hoc Tukey’s tests on linear model (LM), p < 0.05 for all comparisons) (Figure 2, Table 1, Supplementary Table S4). However, these patterns varied across donors (liquid media LM hostpair:plasmid interaction F(2,87) = 3.7, p < 0.029; solid media host:plasmid interaction F(1,28) = 18.3, p < 0.001), such that in experiments with isogenic donors (*S. sonnei* 216 donation to *E. coli*) we detected no difference between the plasmids in liquid media (post-hoc Tukey’s test p = 1) and increased conjugation rate of pKSR100 versus pAPR100 on solid media (p = 0.025). This indicates that the conjugation efficiency differences between the plasmids observed with the native host donors may be due to the different donor backgrounds (Supplementary Table S4). In support of the importance of donor effects, the isogenic *S. sonnei* donor facilitated a greater conjugation efficiency than the native *S. flexneri* donors (all relevant post-hoc comparison p < 0.05) (Supplementary Table S4). Recipient effects were also important as the *E. coli* recipient facilitated more efficient conjugation than *S. sonnei* (all relevant post-hoc comparison p < 0.05) (Supplementary Table S4).

**Table 1.** The impact of plasmid, host, media, and time variables on conjugation efficiencies (ANOVA output of linear models)

| Media | Variables | Df | Sum sq | Mean Sq | F value | p-value |  |
| --- | --- | --- | --- | --- | --- | --- | --- |
| Liquid | hostpair | 2 | 0.5799 | 0.289952 | 182.7608 | <2.20E-16 | *** |
|  | Plasmid | 1 | 0.02199 | 0.021988 | 13.8595 | 0.000349 | *** |
|  | Minutes | 1 | 0.00026 | 0.000263 | 0.1655 | 0.685175 |  |
|  | hostpair:Plasmid | 2 | 0.01173 | 0.005864 | 3.6959 | 0.028803 | * |
|  | hostpair:Minutes | 2 | 0.01441 | 0.007207 | 4.5429 | 0.013287 | * |
|  | Residuals | 87 | 0.13803 | 0.001587 |  |  |  |
| Solid | hostpair | 1 | 2.2918 | 2.29182 | 15.7338 | 0.00046 | *** |
|  | Strain | 1 | 0 | 0.00002 | 0.0002 | 0.989826 |  |
|  | hostpair:Strain | 1 | 2.6617 | 2.66167 | 18.2729 | 0.0002 | *** |
|  | Residuals | 28 | 4.0785 | 0.14566 |  |  |  |

Different levels of conjugation seen across time were also observed with the native donors in liquid media, with a trend of decreasing conjugation rates through time (Supplementary Figure S2), though again this varied with donor and recipient (time:hostpair interaction, F(2,87) = 4.5, p < 0.015) with conjugations into *S. sonnei* displaying increased conjugation rates over time (post-hoc comparisons with other host-pairs p < 0.05). There were also different conjugation dynamics between liquid and solid media, suggesting that conjugation environment also affects conjugation efficiency (solid media: R^2^ = 0.50, F(3,28) = 11.34, p<0.000) (Supplementary Table S4, Figure 2). In general, conjugation was found to be less efficient on solid media than in liquid media (native donors: liquid median = -10.70 logCE, solid median = - 13.04 logCE, Mann-Whitney U = 509, p = 0.0000; isogenic donors: liquid median = - 9.79 logCE, solid mean = -12.43 logCE, Mann-Whitney U = 512, p = 0.0000). Together, these results showed that pAPR100 had a higher conjugation efficiency than pKSR100 from the native hosts, but that this comparative relationship varied with donor and recipient strains and environmental conditions.

### pKSR100 has little fitness cost and alleviates the SOS response

As plasmid burden on bacterial fitness is commonly investigated as a factor contributing to plasmid population dynamics, we investigated the fitness cost of plasmid carriage during growth. Comparing the growth of plasmid-bearing and plasmid-free strains revealed that pAPR100 imposed ∼20% fitness cost in *E. coli* MG1655 whereas pKSR100 did not have a significant fitness cost (Figure 3A). Similar patterns were observed in *S. sonnei* 216, though unlike in *E. coli* we observed a small (∼6%) cost of pKSR100 in *S. sonnei*, suggesting that host factors also contribute to the overall relative fitness impact of plasmid carriage (Figure 3A). Collectively, pAPR100 imposed an increased fitness cost relative to pKSR100 across the hosts.

**Figure 3.**
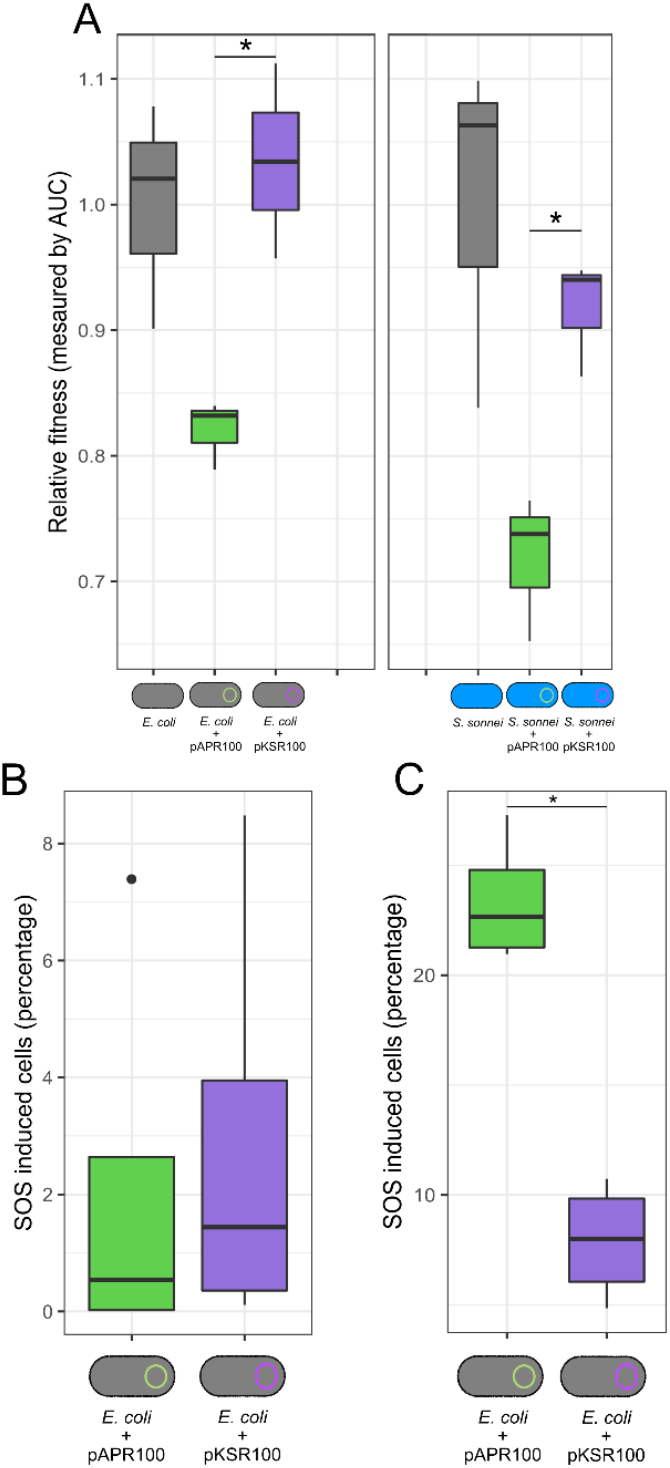
Relative fitness cost and SOS response induction during conjugation and antimicrobial exposure of pKSR100 and pAPR100. (A) Relative fitness of *E. coli* MG1655 (grey icons) and *S. sonnei* 216 (blue icons) carrying either pKSR100 (purple) or pAPR100 (green) compared to plasmid-free wild type (grey bars in the graph). Asterisks denote significance where p=0.01071 for *E. coli* strains and p=0.009971 for *S. sonnei* strains as determined by two sample t-test. (B and C) SOS response levels by plasmid as a proportion of cells in which SOS is induced (measured by GFP expression using a reporter plasmid, see methods) during conjugation (B) and following a two-hour exposure to sub-inhibitory concentrations of ciprofloxacin (C). Individual box plots represent the median, range, and IQR of four independent biological replicate data points adjusted to a negative control for each replicate (see methods, Supplementary Figure 3). No statistically significant difference was observed between the two plasmids during conjugation (p=0.7864), but there was a marked level of SOS response alleviation in cells carrying pKSR100 (p=0.000237) as determined by a two sample t-test.

The comparative genomic analysis revealed that pKSR100 contained a cluster of five genes including single stranded DNA binding protein gene, plasmid SOS inhibition A (*psiA*) and B *(psiB*) genes which were absent in pAPR100 (Figure 1C, Figure S1A) and are known to alleviate the bacterial SOS response in *E. coli* (34). A gene encoding for a hypothetical protein and an uncharacterised protein family (UPF) 0401 protein *ykfF* gene was also found in the same cluster though their roles in SOS response alleviation, if any, is unclear. The SOS response is often induced during conjugation and by environmental stressors and is known to be costly (35). Given the disparate fitness costs between pKSR100 and pAPR100, we investigated whether there were differences in SOS response and alleviation attributable to the plasmids. Firstly, because of the differences between conjugation genes and the known activation of the SOS response during conjugation (35), we measured SOS response induction during conjugation anticipating that pAPR100 conjugation might elicit a greater SOS response in the recipients due to the absence of the gene cluster containing *psiB*, and that pKSR100 (which has the *psiB* gene cluster) would alleviate the SOS response. However, we observed no statistically significant difference among SOS induction between the plasmids during conjugation, though both showed a slight increase in SOS relative to the negative control (Figure 3B).

Because these plasmids derive from a patient community with high levels of antimicrobial use, we then sought to determine if either plasmid alleviated the SOS induction in response to sublethal concentrations of antibiotics. Specifically, we investigated the fluoroquinolone antibiotic ciprofloxacin, which is: known to induce bacterial SOS response as a result of DNA mutagenesis stimulation in *E. coli* (36); the recommended, and overwhelmingly administered, treatment for shigellosis in this patient community (37, 38); and an antimicrobial to which neither pKSR100 or pAPR100 confer resistance (see below and Table 2). Our subsequent measuring of SOS induction among cells (carrying either pKSR100 or pAPR100) after a two-hour exposure to subinhibitory concentrations of ciprofloxacin revealed that cells harbouring pKSR100 were significantly better at alleviating the SOS induction than cells carrying pAPR100 (Figure 3C). Thus cells harbouring pKSR100 were less likely to activate the costly SOS response following exposure of sublethal concentrations of an antimicrobial for which the plasmid conferred no resistance.

**Table 2.**
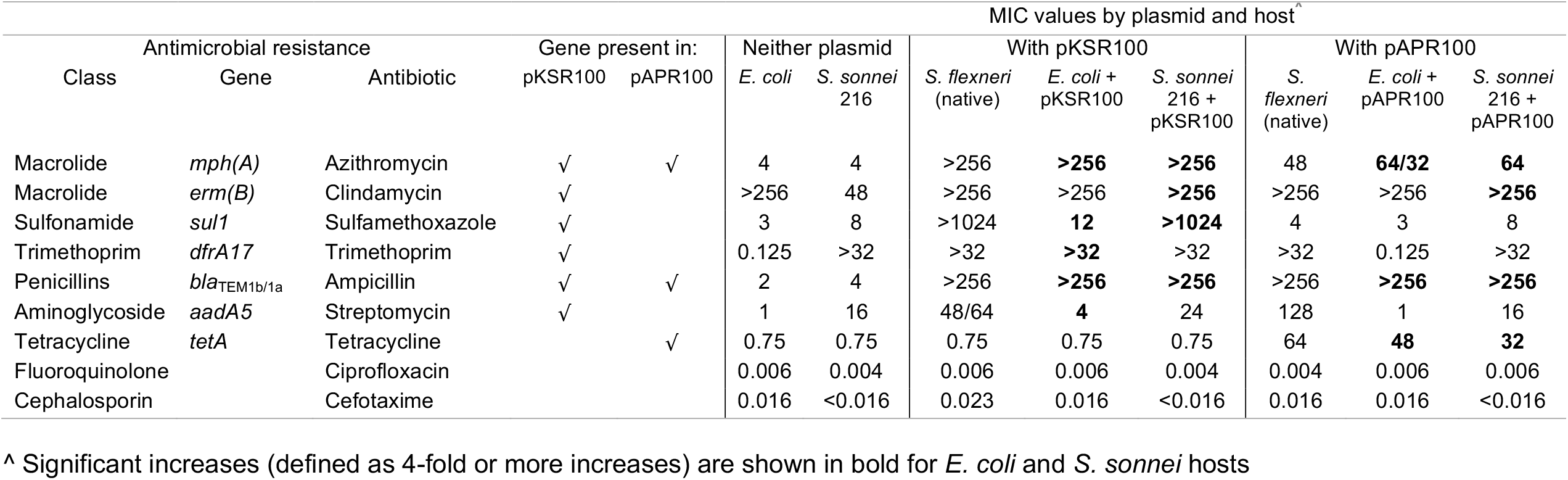
The impact of pKSR100 and pAPR100 on antimicrobial resistance phenotypes

### pKSR100 offers a greater range and magnitude of AMR

Owing to the importance of AMR in driving emergence of shigellosis among MSM populations, we also investigated the AMR phenotypes conferred by the two plasmids. Predictions of AMR genes identified six AMR genes in pKSR100, which accounted for 50% (6 of 12) of the total AMR genes in the host strain of *S. flexneri* 2a, while pAPR100 only had three AMR genes contributing 27% (3 of 11) to the total AMR gene content of its host *S. flexneri* 2a strain (Supplementary Table S2 and S3). Analysis of AMR genes present on the plasmids using ResFinder predicted that pKSR100 encoded resistance to sulphonamides, macrolides, trimethoprim, beta-lactams, and aminoglycosides and that pAPR100 encoded resistance to macrolides, beta-lactams, and tetracycline showing that the AMR genes of the plasmids differed both in the number and variety of drug classes they provided resistance against (Table 2). Thus, both resistance plasmids conferred multidrug resistance and contained resistance genes relevant to transmission among MSM, but pKSR100 was predicted to confer broader-spectrum resistance.

To test the genotypic predictions of increased AMR conferred by the plasmids, we correlated our genotypic information with minimum inhibitory concentration (MIC) phenotypes. The antimicrobials tested were based on the genotypic AMR predictions, and to differentiate plasmid and host factors, MICs were determined for the native hosts, and in both *E. coli* and *S. sonnei* 216 transconjugants of both plasmids compared to plasmid-free wildtypes (Table 2). The two plasmid-free hosts carried intrinsic resistance to three antibiotics *i*.*e. E. coli* was resistant against clindamycin and *S. sonnei* 216 to trimethoprim and streptomycin. Collectively however, the results demonstrated general agreement between the AMR genotype carried on plasmids and phenotype with a minimum four-fold increase in the MIC values compared to the controls when the AMR genes were present (Table 2). Notably however, pKSR100 conferred a higher level of resistance to azithromycin, a phenotype critical for driving circulation of shigellosis in this community at that time, by virtue of encoding the additional macrolide resistance gene *ermB* (Table 2, (23, 24)).

In addition to those antimicrobials where resistance was anticipated to change based on genotypic prediction, we tested whether the plasmids conferred AMR against other clinically relevant antimicrobials; ciprofloxacin and cefotaxime. This was to ensure we were not missing relevant AMR conferred by potentially novel genes, or potentiating genes, and to ensure the robustness of our SOS induction results. As expected, neither plasmid altered the MIC values against these antimicrobials.

Collectively, these AMR results show that pKSR100 confers resistance to a more extensive range of antimicrobials, and a higher level of resistance to azithromycin, than pAPR100.

## Discussion

Here, we investigated two AMR plasmids with different epidemiological outcomes from the same clinical niche to identify putative plasmid phenotypes associated with epidemiological success, using comparative genomics as a guide. The spread and persistence of plasmids may depend on several plasmid and host associated factors such as the plasmid type, host range, conjugative capacity, and fitness cost (39-44). Understanding the contributions of each of these factors is important in determining the fate of AMR plasmids found in clinical settings and predicting evolutionary trajectories of those plasmids (16). Our results identified that the epidemiologically successful pKSR100 conferred increased AMR, reduced fitness cost, and reduced SOS response in the presence of subinhibitory concentrations of antibiotics, compared with the less successful pAPR100, despite pKSR100 having a lower conjugation efficiency in native hosts.

The epidemiologically prominent plasmid, pKSR100, carried more AMR genes than pAPR100. In addition to the AMR genes carried on the plasmid, the bacterial host also carried AMR genes on the chromosome (Tables S2, S3). However, there were few discrepancies between the non-plasmid AMR gene content of the native hosts (only a *dfrA1* and additional *bla*TEM gene), and these did not result in phenotypic differences between the native hosts against the cognate antimicrobials (trimethoprim and ampicillin respectively, Table 2). Thus, it is unlikely that there was a relationship between the AMR genes present on the respective bacterial chromosomes (*i*.*e*. the *S. flexneri* 2a major and minor sublineages) and the spread of the plasmids. This is further evidenced by the remarkable onward global spread of pKSR100 from the setting in which it was originally described (Figure 1). Thus, it is possible that the more extensive range of AMR and higher azithromycin resistance conferred by pKSR100 relative to pAPR100 was a key determining factor in the resulting epidemiological success and widespread dissemination of pKSR100. However, given that AMR genes frequently recombine between plasmids, and both plasmids already clearly encoded MDR, we are still faced with the question of why it was pKSR100 that acquired the broader AMR profile and went on to globally disseminate despite the comparative longevity of circulation of pAPR100 in the community. To address this, we considered further non-AMR factors guided by the comparative genomics, namely conjugation rate and fitness cost.

Conjugation is an efficient mechanism for plasmid dissemination and has been widely studied (1). Theoretically, a higher conjugative capacity might result in a greater spread of plasmids. However, in our case, we observed that the globally widespread pKSR100 had a lower conjugation efficiency than pAPR100. The conjugative ability of plasmids can depend on donor or recipient strains’ background as well as abiotic environmental factors, but these were not found to overcome the generality in our system that pKSR100 was less conjugative or equivocal than pAPR100 from their respective native hosts (Figure 2). The lack of apparent importance of the conjugation efficiency exclusively for broader spread is consistent with a previous study examining the contribution of conjugation and other factors such as fitness effects towards plasmid spread in hospital settings (12). However, we also observed that the differences of conjugation efficiency between the two plasmids diminished from an isogenic donor background, suggesting that donor factors are influencing conjugation efficiency in our model. This is not unexpected as the effect of donor and recipient backgrounds are known to affect the conjugative capacity of plasmids (45). Hence, our results are consistent with previous studies indicating that conjugation efficiency is dependent on the donor, recipient, and environment, and may not be a major contributor to dissemination among bacterial populations in real world settings.

By contrast, the lack of a fitness cost imposed by pKSR100 may have been an important factor in perpetuating its epidemiological success. The fitness costs imposed by plasmids depend on the plasmid and host combination as well as their environmental factors along with gene conflicts (39, 46) and, previous work has demonstrated that fitness cost measured in a laboratory setting might not translate to success in the natural environment (22). We addressed host variability by measuring the fitness cost imposed by our plasmids in multiple backgrounds, and the broader database analysis (Figure 1, S1B) indicated a similar host range for the two plasmids and their close relatives. While our results may not relate exactly to the fitness cost imposed by these plasmids on their natural hosts, it helps us understand the effects that fitness costs may have on transmission through other residents in their natural gut microbe community. Our observation of the unsuccessful pAPR100 having a greater fitness cost is consistent with fitness costs affecting plasmid fates in microbial populations, and suggests that fundamental plasmid properties can contribute to epidemiological outcomes, specifically in a community-transmitting pathogen in our case.

Given the difference in SOS alleviation gene content between the two plasmids, we also examined the impact of the two plasmids on the induction of the costly SOS response. Since plasmids move as single stranded DNA molecules to recipient cells during conjugation, the possible induction of the SOS response may act as an influencing factor for the dissemination of plasmids. The pKSR100-related plasmid R100 had been shown to induce the SOS response when conjugated into *Vibrio cholerae* recipient (alleviated by the *psiB* gene cluster (47, 48)). Hence, finding no difference in SOS induction during conjugation with the *psiB* cluster-containing pKSR100 and pAPR100 (which does not contain the cluster) in *E. coli*, indicates that SOS inductions by plasmids is host dependent. However, as sub-inhibitory concentrations of ciprofloxacin have been demonstrated to induce the SOS response (49) and this is a highly relevant exposure for our system (as ciprofloxacin is a recommended treatment for shigellosis and other STIs circulating among MSM (37)) we explored the possibility that the pKSR100 may alleviate activation of the costly SOS response during exposure to subinhibitory concentrations of antimicrobials. Our finding that pKSR100 protected against the induction of the SOS response during exposure to subinhibitory concentrations of a clinically relevant antimicrobial supports the notion that an SOS response alleviation phenotype may contribute to AMR emergence in an epidemiological setting of high antimicrobial use.

The results of our study are consistent with the theory that, alongside AMR, plasmid associated non-AMR phenotypes may play a crucial role in facilitating the dissemination of AMR plasmids. Specifically, an epidemiologically successful plasmid conferred increased AMR and alleviated the SOS response at a much-reduced fitness cost compared with an unsuccessful plasmid in the same ecological system/niche. Although our study used only two plasmids from a single model system, the plasmids we used were important examples drawn from an established pathogen and real world epidemiological scenario (capturing complex networks in their natural habitats). Thus, our findings may help guide future studies towards the development of universal ground truths regarding important plasmid phenotypes for AMR emergence and establishment within the dynamic and problematic *Enterobacteriaceae*. We also mitigated the possibility that phenotypic findings may have been dependent on non-plasmid factors by varying the host and environmental conditions, which will vary in the natural habitat. Thus, working backwards from an established epidemiological scenario is a valuable approach for identifying plasmid-associated phenotypes that contribute to the epidemiological trajectory of AMR, and perhaps one day could be turned prospectively to aid prediction and prevention of AMR emergence in future surveillance.

## Materials and methods

### Plasmid comparison and comparative genomics

Full plasmid sequences were extracted from genome sequences from a previous study (24). Specifically, for pKSR100 the isolate corresponding to accession number ERR1364116 was used and for pAPR100 the contiguous sequence 24 from the isolate corresponding with accession number ERR1364014 was used.

Extracted genomic sequences of the plasmids were annotated using RAST server (50) using the default settings. The annotated files were used to generate a BLAST comparison using EasyFig (51). Regions of similarity were identified between the two plasmids using a BLAST cut off of over 95%. Examination of the annotated functions of genes in dissimilar regions using Roary v3.11.2 (52) (Supplementary file S2) showed that these belonged to three main functional groups which were then used as a basis for further experimentation. AMR gene content of the strains and two plasmid sequences used was predicted using ResFinder version 4.1 (53) using default parameters.

Comparison of the plasmid sequences against publicly available data was done in two ways. 1) a BLAST search of the NCBI non-redundant database on 17th May 2021 and retrieval of papers and metadata associated with closely related sequences, and 2) screening against the 661K COBS data structure. The two plasmid sequences were used to query against the COBS index of 661,405 curated draft genomes (33) with a baseline kmer similarity cutoff of 0.4. Characterisation data of genomes containing the sequence of interest was extracted from figshare (https://dx.doi.org/10.6084/m9.figshare.14061752) (33). Additional filtering of hits based on a higher kmer similarity threshold (>= 0.8), a major species abundance of at least 90% and if the genome was determined to be of high quality (as defined in (33)) was applied. ggplot2 v3.3.5 was used to visualise the 661K COBS search results and Inkspace v1.1 was used to manually edit plots where needed.

### Bacterial conjugation

All bacterial strains used in conjugation experiments were grown in TSB overnight and diluted 1:100 into fresh media and grown for three hours before preparing the conjugation mixture. Donor and recipient strains were mixed 1:1 in a final volume of 500µL for the conjugation experiments in liquid media and incubated shaking at 215 rpm at 37°C for 75 minutes. Conjugation mixtures were serially diluted and plated on selective media to distinguish between donors, recipients, and transconjugants every 15 minutes and the resulting colony forming units were counted to enumerate each. Conjugation mixtures were prepared as described above for the conjugation experiments on solid media and 10µL of the conjugation mixture was spotted onto a sterile nitrocellulose filter paper placed on an agar plate and incubated at 37°C without shaking for five hours. Conjugation mixtures present on the filter papers were submerged in 500µL of sterile PBS, agitated by vortexing, serially diluted, and plated as above every hour for CFU measurements. Conjugation efficiencies (CE) for four biological replicates were calculated as previously described (54) using the following equation:

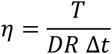

Where η = conjugation efficiency (CE) measured in mL cell^-1^ h^-1^, T = transconjugants, D = donors, R = recipients and Δt = total conjugation time.

### Linear Modelling of conjugation

Linear models (LM) were created to examine the conjugation efficiency of the plasmids while accounting for the effects of donor and recipient strain backgrounds. Linear modelling was performed using R(v4.0.4), and the ResourceSelection (v0.3-5), ggplot2 (v3.3.4)), ggfortify (V0.4.13), stats (v4.1.0), car (v3.0-11), nortest(1.0-4) and MASS (v7.3-54) packages. Conjugation was modelled separately for solid and liquid media due to differences in sampling times, and time points lacking data for one or more replicates (due to lack of transconjugants) were excluded. For each media type we fitted a full model (with plasmid, host pair, time, and all interactions) and performed model reduction by AIC comparisons between nested models.

Host pair was either 1) *S. flexneri* 2a (heterogeneous native donors) to *E. coli* recipient, 2) *S. sonnei* 216 (isogenic donor) to *E. coli* recipient, or 3) *S. flexneri* 2a (heterogeneous native donors) to *S. sonnei* 216. The data for the liquid media conjugation model was –log_e_ transformed to meet assumptions of linear modelling. Pairwise contrasts were estimated for each significant combination of factors using emmeans (1.7.1-1) (Table 1).

#### SOS response measurements during conjugation

SOS induction during conjugation was measured by using GFP expression as a proxy by means of a Pint-gfp fusion reporter plasmid p9092 (a kind gift from Didier Mazel (55)). Expression of the integrase promoter Pint is a strong signal of SOS induction (55). Briefly, p9092 was transformed into *E. coli* MG1655 that constitutively expressed mCherry and was used as the recipient strain. Donor strains were *E. coli* MG1655 strains carrying either pKSR100 or pAPR100 with no fluorescence protein expression and *E. coli* MG1655 with no plasmids was used as the negative control for conjugation. Samples were set up as with the conjugation experiments and after one hour, GFP expressing cells in the samples were counted using a Bio-Rad ZE5 Cell Analyzer flow cytometer with a total count of 100,000 for each sample. The resulting data files were analysed using FCS Express version 7 (De Novo Software) to determine the percentage of GFP expressing cells in the mCherry population of the sample. Experiments were carried out with four independent biological replicates.

#### SOS response measurements during ciprofloxacin exposure

*E. coli* MG1655 with no fluorescence protein expression carrying p9092 was transformed with either pKSR100 or pAPR100 and the presence of plasmids were confirmed by growth on both tetracycline (selection for p9092) and azithromycin (selection for pKSR100 and pAPR100). Individual colonies of these strains were grown overnight at 37°C with shaking in a 96 well plate in TSB supplemented with 15µg/mL tetracycline and sub-cultured 1:100 into fresh TSB again with no antibiotics in a 96 well plate and grown in similar conditions for 2-3 hours to have the cultures in exponential growth phase. These cultures were then challenged with 0.0025 µg/mL ciprofloxacin (1/4 MIC) for 2 hours before quantifying the GFP expressing cells using a Bio-Rad ZE5 Cell Analyzer. The percentage of GFP expressing cells were determined using the cells not exposed to ciprofloxacin as a negative control using FCS Express version 7 (De Novo Software). Four individual biological replicates were tested for each plasmid.

#### Bacterial growth curves and fitness

Transconjugant *E. coli* strains (with constitutive GFP expression) or *S. sonnei* 216 strains, carrying pKSR100 or pAPR100 from conjugation experiments were frozen in 25% glycerol and used as the source freezer stock for the growth curves. Pre-cultures for the bacterial growth curves were grown in M9 medium without azithromycin selection overnight incubating at 37°C with 215 rpm shaking and diluted 1:100 into fresh M9 media adjusting all the cultures to the lowest OD600 value. The cultures were then distributed into a sterile 96-well plate (Greiner Bio One, UK) with a moisture barrier seal (4titude, UK) and incubated at 37°C with shaking in a Synergy H1 multi-mode plate reader taking optical density readings at 600nm every 15 minutes. The resulting values were plotted using the R package Growthcurver (56) to determine the area under the curve (AUC) for each growth curve. Average values of the technical replicates for AUC of each of the plasmid carrying strains were normalised using the average AUC value of the technical replicates of plasmid free *E. coli* and *S. sonnei* 216 strains to obtain relative fitness of each of the transconjugant *E. coli* and *S. sonnei* 216 strains for three biological replicates.

### Minimum Inhibitory Concentration measurements

MIC measurements were carried out using Liofilchem® MIC test strips (Liofilchem, Italy) following manufacturer’s guidelines. Bacterial inocula for the MIC testing were prepared following the EUCAST guidelines for broth microdilution testing breakpoint table (https://www.eucast.org/fileadmin/src/media/PDFs/EUCAST_files/Breakpoint_tables/v_11.0_Breakpoint_Tables.pdf) and were spread on Mueller Hinton Agar plates (Bio-Rad, France) using sterile cotton swabs after which the MIC test strip was applied, and plates were incubated at 37°C for 18 hours before the readings were recorded.

## Acknowledgements

This work was supported by an Academy of Medical Sciences Springboard award (SBF002\1114). KSB was supported by a Wellcome Trust Clinical Research Career Development Award (106690/A/14/Z). GES is supported by a studentship from the MRC Discovery Medicine North (DiMeN) Doctoral Training Partnership (MR/N013840/1). MDS is supported by a Biotechnology and Biological Sciences Research Council project grant BB/V009184/1. RJB is supported by a Medical Research Council and UK Department for International Development (DFID) project grant MR/R020787/1. Kate Baker is affiliated to the National Institute for Health Research Health Protection Research Unit (NIHR HPRU) in Gastrointestinal Infections at University of Liverpool in partnership with UK Health Security Agency (UKHSA), in collaboration with University of Warwick. The views expressed are those of the author(s) and not necessarily those of the NHS, the NIHR, the Department of Health and Social Care or UKHSA. We would like to thank Michael Bottery (University of Manchester) for the kind gift of GFP and mCherry tagged *E. coli* MG1655 strains and Didier Mazel for his kind gift of the Pint-gfp fusion reporter plasmid p9092.

## Supplementary Materials

**Table S1.** Strains and plasmids used in this study

| Strain/Plasmid | Description | Reference |
| --- | --- | --- |
| *Shigella flexneri* 2a | (Native strains) |  |
| ERR1364116 | Major sublineage clinical strain naturally carrying pKSR100 | Baker *et al* 2018 |
| ERR1364014 | Minor sublineage clinical strain naturally carrying pAPR100 | Baker *et al* 2018 |
| *Shigella sonnei* |  |  |
| ERR1364216 | Clinical strain naturally not carrying either pKSR100 or pAPR100 (referred to as *S. sonnei* 216 in the main text) | Baker *et al* 2018 |
| *Escherichia coli* |  |  |
| MG1655 | Laboratory strain |  |
| MG1655::GFP | Constitutive GFP expressing *E. coli* MG1655 strain | Bottery lab |
| MG1655:mCherry | Constitutive mCherry expressing *E. coli* MG1655 strain | Bottery lab |
| Plasmids |  |  |
| pKSR100 | Azithromycin resistance plasmid from *S. flexneri* 2a ERR1364116 | Baker *et al* 2018 |
| pAPR100 | Azithromycin resistance plasmid from *S. flexneri* 2a ERR1364014 | Baker *et al* 2018 |
| p9092 | Reporter plasmid with *Pint-gfp* fusion | Mazel Lab |

**Table S2.** ResFinder results for pKSR100 carrying strain ERR1364116 with AMR genes associated with pKSR100 in bold

| Resistance gene | Identity | Alignment Length/Gen e Length | Coverage | Position in reference | Contig | Position in contig | Phenotype | Accession no. |
| --- | --- | --- | --- | --- | --- | --- | --- | --- |
| ***sul1*** | **100** | **840/840** | **100** | **1..840** | **\|SC\|contig000011** | **23726..24565** | **Sulphonamide resistance** | **U12338** |
| *catA1* | 99.85 | 660/660 | 100 | 1..660 | \|SC\|contig000010 | 70367..71026 | Phenicol resistance | V00622 |
| *mdf(A)* | 98.22 | 1233/1233 | 99.513382 | 1..1233 | \|SC\|contig000008 | 496420..497646 | Warning: gene is missing from Notes file. Please inform curator. | Y08743 |
| ***mph(A)*** | **100** | **906/906** | **100** | **1..906** | **\|SC\|contig000011** | **29876..30781** | **Macrolide resistance** | **D16251** |
| ***erm(B)*** | **99.86** | **738/738** | **100** | **1..738** | **\|SC\|contig000011** | **31963..32700** | **Macrolide resistance** | **JN899585** |
| ***dfrA17*** | **100** | **474/474** | **100** | **1..474** | **\|SC\|contig000011** | **21787..22260** | **Trimethoprim resistance** | **FJ460238** |
| *blaOXA-1* | 100 | 831/831 | 100 | 1..831 | \|SC\|contig000010 | 77101..77931 | Beta-lactam resistance | HQ170510 |
| ***bla<sub>TEM-1B</sub>*** | **100** | **861/861** | **100** | **1..861** | **\|SC\|contig000011** | **18001..18861** | **Beta-lactam resistance**<br>**Alternate name; RblaTEM-1** | **AY458016** |
| ***aadA5*** | **100** | **789/789** | **100** | **1..789** | **\|SC\|contig000011** | **22391..23179** | **Aminoglycoside resistance** | **AF137361** |
| *aadA1* | 99.75 | 792/792 | 100 | 1..792 | \|SC\|contig000010 | 78044..78835 | Aminoglycoside resistance | JQ414041 |
| *tet(B)* | 100 | 1206/1206 | 100 | 1..1206 | \|SC\|contig000010 | 66170..67375 | Tetracycline resistance | AF326777 |

**Table S3.** ResFinder results for pAPR100 carrying strain ERR1364014 with AMR genes associated with pAPR100 in bold

| Resistance gene | Identity | Alignment Length/Gene Length | Coverage | Position in reference | Contig | Position in contig | Phenotype | Accession no. |
| --- | --- | --- | --- | --- | --- | --- | --- | --- |
| *aadA1* | 99.75 | 792/792 | 100 | 1..792 | \|SC\|contig000005 | 26837..27628 | Aminoglycoside resistance | JQ414041 |
| *aadA1* | 100 | 789/789 | 100 | 1..789 | \|SC\|contig000021 | 930788..931576 | Aminoglycoside resistance | JQ480156 |
| *catA1* | 99.85 | 660/660 | 100 | 1..660 | \|SC\|contig000005 | 19159..19818 | Phenicol resistance | V00622 |
| *mdf(A)* | 98.22 | 1233/1233 | 99.513382 | 1..1233 | \|SC\|contig000020 | 250941..252167 | Warning: gene is missing from Notes file. Please inform curator. | Y08743 |
| ***mph(A)*** | **100** | **906/906** | **100** | **1..906** | **\|SC\|contig000024** | **44542..45447** | **Macrolide resistance** | **D16251** |
| ***tet(A)*** | **100** | **1200/1200** | **100** | **1..1200** | **\|SC\|contig000024** | **35010..36209** | **Tetracycline resistance** | **AJ517790** |
| *tet(B)* | 100 | 1206/1206 | 100 | 1..1206 | \|SC\|contig000005 | 29549..30754 | Tetracycline resistance | AF326777 |
| *dfrA1* | 100 | 474/474 | 100 | 1..474 | \|SC\|contig000021 | 932253..932726 | Trimethoprim resistance | X00926 |
| *blaOXA-1* | 100 | 831/831 | 100 | 1..831 | \|SC\|contig000005 | 25894..26724 | Beta-lactam resistance | HQ170510 |
| ***bla<sub>TEM-1A</sub>*** | **100** | **861/861** | **100** | **1..861** | **\|SC\|contig000024** | **33049..33909** | **Beta-lactam resistance Alternate name; RblaTEM-1** | **HM749966** |
| *bla<sub>TEM-1A</sub>* | 99.88 | 861/861 | 99.88385598 | 1..861 | \|SC\|contig000011 | 12425..13284 | Beta-lactam resistance Alternate name; RblaTEM-1 | HM749966 |

**Figure S1A.**
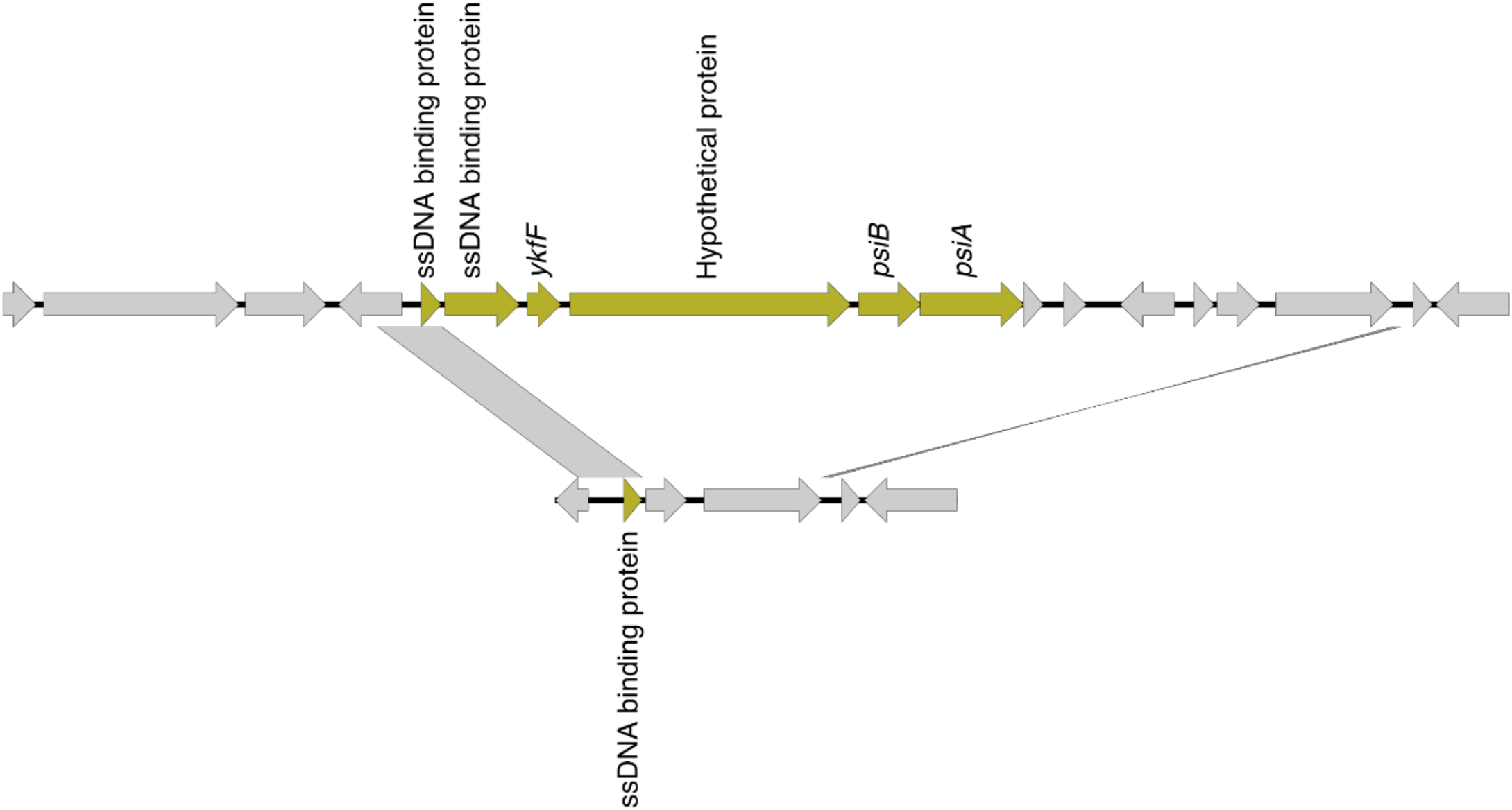
Zoomed in version of the SOS response alleviation gene cluster (green) in pKSR100 (top), which is absent in pAPR100 (bottom). NB: The “Hypothetical protein” in pKSR100 is related to *parB* based on BLASTn searching.

**Figure S1B.**
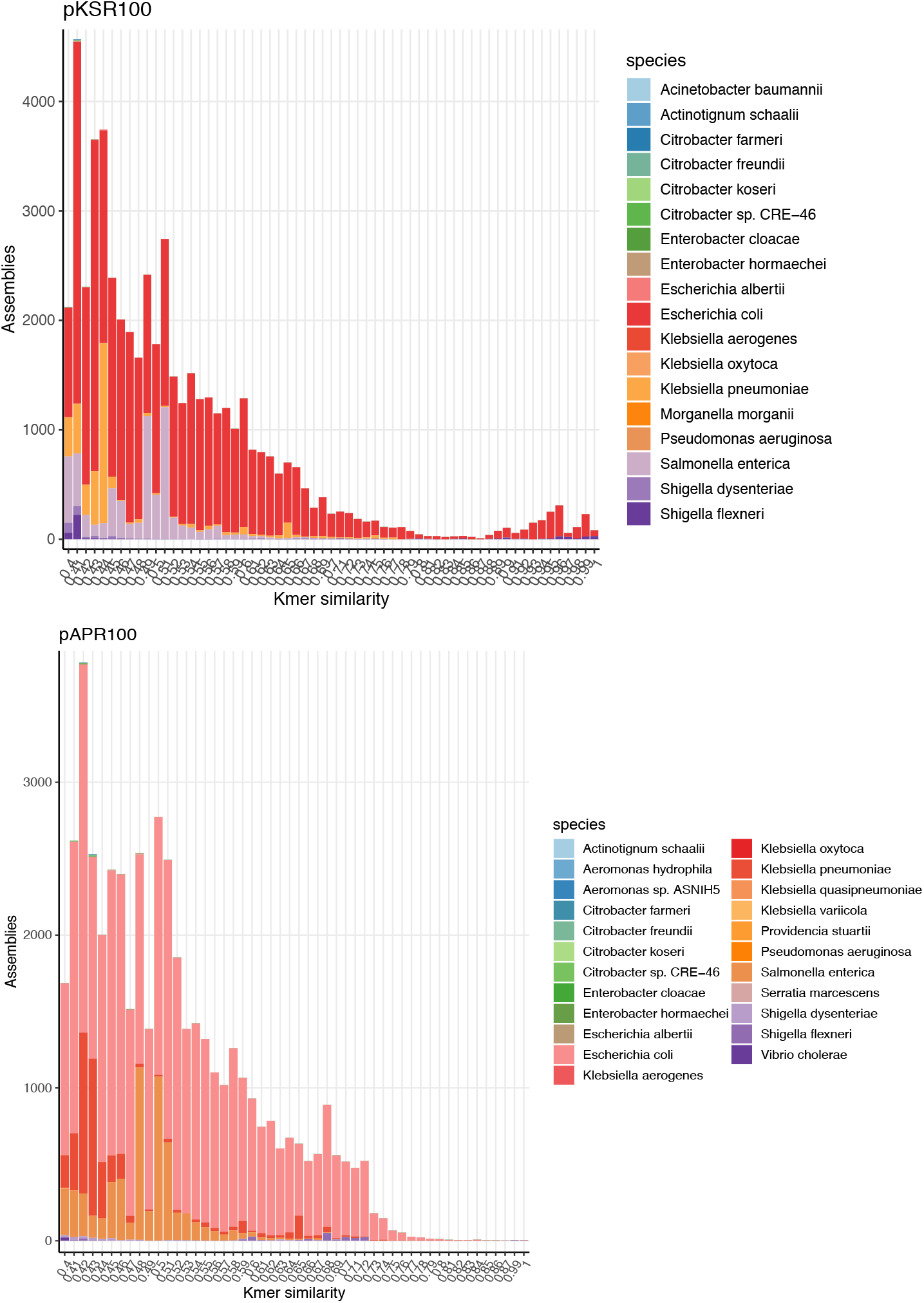
Species distribution containing more distant (down to 0.4 kmer similarity) relatives of pKSR100 (upper) and pAPR100 (lower) in the COBS661K data structure. In both cases, the dominant host species are *E. coli, S. enteritica*, and *K. pneumoniae* coloured according to each inlaid key.

**Figure S2.**
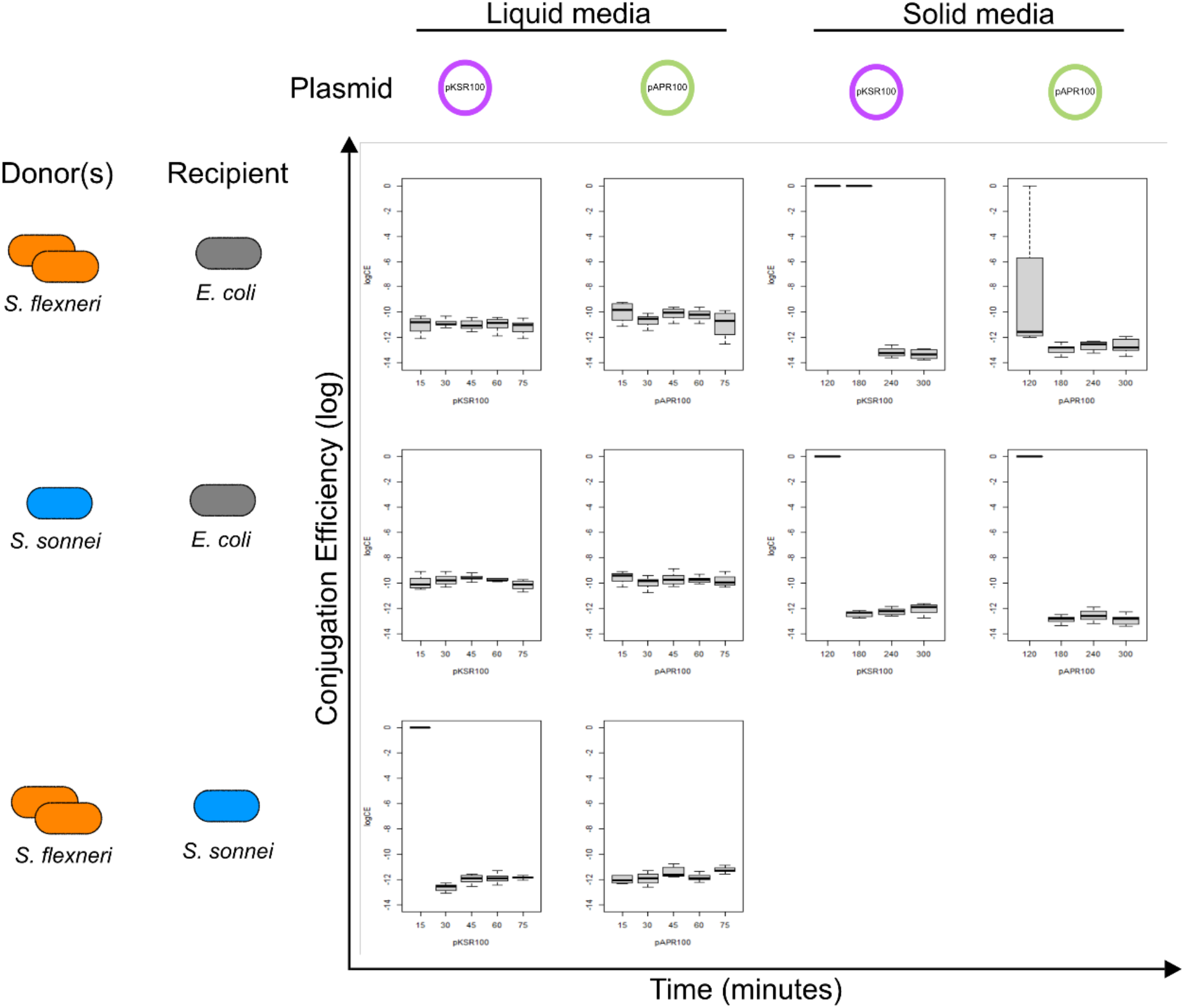
Full dataset (including time) used for linear models (See Figure 2, Table 1 and Table S4). Time (minutes) is shown on the x-axis and conjugation efficiencies on the y-axis. The graphs are arranged in the grid by donor/recipient/plasmid/media combinations as indicated by inlaid keys.

**Table S4.** The impact of plasmid, host, media, and time variables on conjugation efficiencies (linear model post-hoc pairwise comparisons output)

| Comparison | Conditions | Estimate | SE | df | t ratio | P-value |  |
| --- | --- | --- | --- | --- | --- | --- | --- |
| pAPR100 - *E. coli* vs *S. sonnei* recipient | Liquid media Native donors | 0.104501 | 0.014082 | 87 | 7.420706 | 1.40E-09 | **** |
| pAPR100 - native donors vs isogenic donor | Liquid media *E. coli* recipient | -0.06451 | 0.014082 | 87 | -4.58096 | 0.000218 | **** |
| pAPR100 vs pKSR100 in native donors | Liquid media *E. coli* recipient | 0.049622 | 0.014082 | 87 | 3.523662 | 0.008643 | ** |
| pAPR100 to *E. coli* vs pKSR100 to *S. sonnei* | Liquid media Native donors | 0.146354 | 0.014082 | 87 | 10.39271 | 3.09E-10 | **** |
| pAPR100 in native donors vs pKSR100 in isogenic donor | Liquid media *E. coli* recipient | -0.06518 | 0.014082 | 87 | -4.62849 | 0.000182 | **** |
| pAPR100 - native donors to *S. sonnei* vs isogenic donor to *E. coli* | Liquid media | -0.16901 | 0.014082 | 87 | -12.0017 | 3.09E-10 | **** |
| pAPR100 to *S. sonnei* recipient vs pKSR100 to *E. coli* recipient | Liquid media Native donors | -0.05488 | 0.014082 | 87 | -3.89704 | 0.002554 | *** |
| pAPR100 vs pKSR100 to *S. sonnei* recipient | Liquid media Native donors | 0.041853 | 0.014082 | 87 | 2.972008 | 0.042927 | * |
| pAPR100 in native donors to *S. sonnei* recipient vs pKSR100 in isogenic donor to *E. coli* recipient | Liquid media | -0.16968 | 0.014082 | 87 | -12.0492 | 3.09E-10 | **** |
| pAPR100 in isogenic donor vs pKSR100 in native donors | Liquid media *E. coli* recipient | 0.114132 | 0.014082 | 87 | 8.104618 | 3.54E-10 | **** |
| pAPR100 isogenic donor to *E. coli* vs pKSR100 native donors to *S. sonnei* | Liquid media | 0.210865 | 0.014082 | 87 | 14.97367 | 3.09E-10 | **** |
| pAPR100 vs pKSR100 in isogenic donor | Liquid media *E. coli* recipient | -0.00067 | 0.014082 | 87 | -0.04754 | 1 |  |
| pKSR100 - *E. coli* vs *S. sonnei* recipient | Liquid media Native donors | 0.096733 | 0.014082 | 87 | 6.869052 | 1.40E-08 | **** |
| pKSR100 - Native donors vs isogenic donor | Liquid media *E. coli* recipient | -0.1148 | 0.014082 | 87 | -8.15216 | 3.45E-10 | **** |
| pKSR100 - native donors to *S. sonnei* vs isogenic donor to *E. coli* | Liquid media | -0.21153 | 0.014082 | 87 | -15.0212 | 3.09E-10 | **** |
| *E. coli* vs *S. sonnei* recipient (native donors) | Conjugation through time | -0.00165 | 0.000594 | 87 | -2.77543 | 0.018336 | * |
| Native vs isogenic donors (*E. coli* recipient) | Conjugation through time | -0.00022 | 0.000594 | 87 | -0.36936 | 0.927624 |  |
| Native donor to *S. sonnei* recipient vs Isogenic donor to *E. coli* recipient | Conjugation through time | 0.001429 | 0.000594 | 87 | 2.406074 | 0.04746 | * |
| pAPR100 - native donors vs isogenic donor | *Solid media*<br>*E. coli* recipient | 0.04157 | 0.190828 | 28 | 0.21786 | 0.996262 |  |
| pAPR100 vs pKSR100 in native donors | *Solid media*<br>*E. coli* recipient | 0.57507 | 0.190828 | 28 | 3.01356 | 0.026301 | * |
| pAPR100 in native donors vs pKSR100 in isogenic donor | *Solid media*<br>*E. coli* recipient | -<br>0.536972 | 0.190828 | 28 | -<br>2.813898 | 0.04153 | * |
| pAPR100 in isogenic donor vs pKSR100 in native donors | *Solid media*<br>*E. coli* recipient | 0.5335 | 0.190828 | 28 | 2.7957 | 0.043255 | * |
| pAPR100 vs pKSR100 in isogenic donor | *Solid media*<br>*E. coli* recipient | -<br>0.578545 | 0.190828 | 28 | -<br>3.031757 | 0.025207 | * |
| pKSR100 - native donors vs isogenic donor | *Solid media*<br>*E. coli* recipient | -<br>1.112045 | 0.190828 | 28 | -5.82746 | 1.66E-05 | **** |

**Figure S3.**
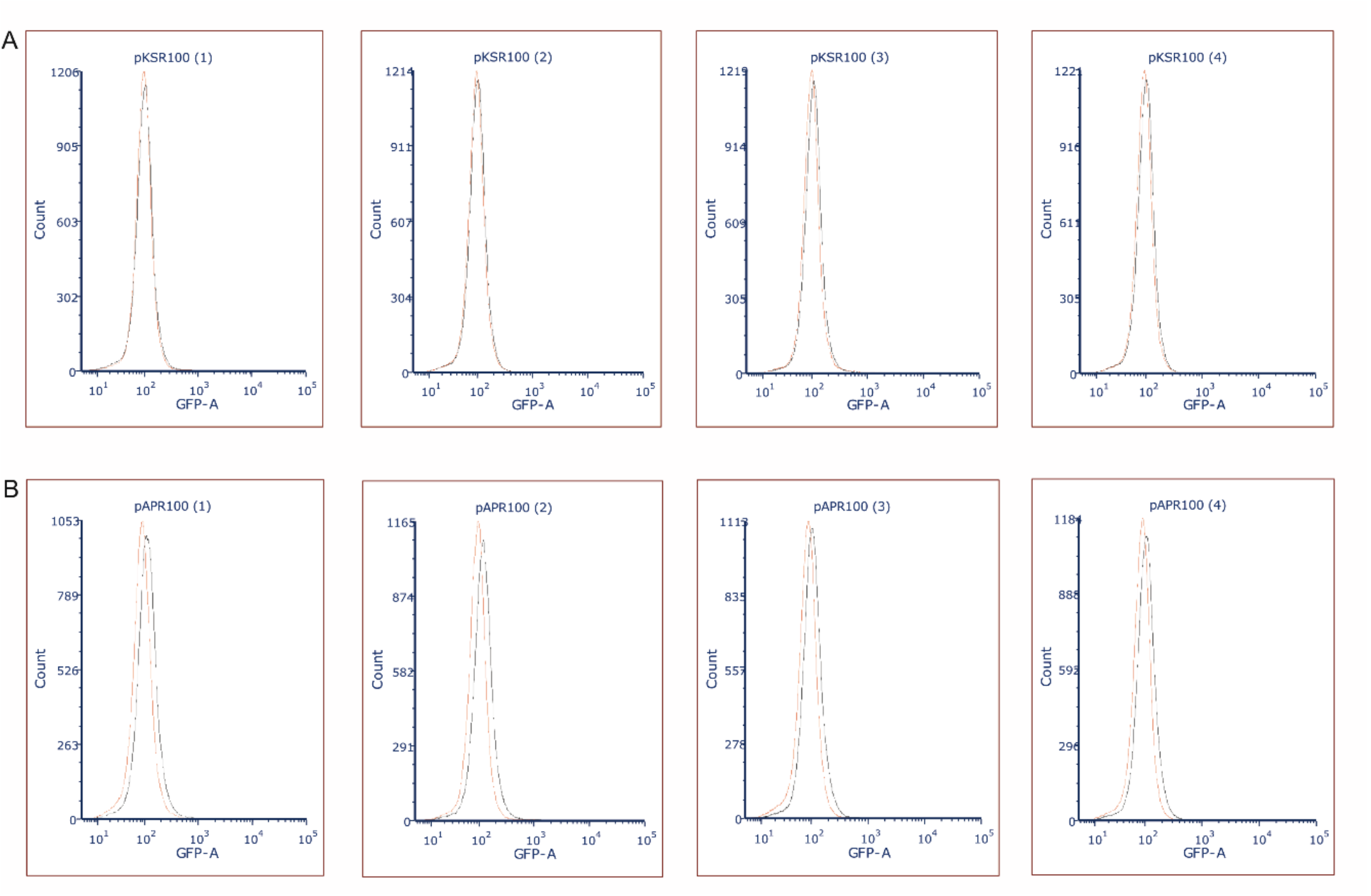
Histograms of SOS response induction (as measured by fluorescence [GFP-A]) by cell count (y-axis) for individual biological replicates represented in Figure 3C showing that (A) pKSR100 was able to alleviate the SOS response (lack of a right shift of black curves) when exposed to ciprofloxacin when compared to the negative control (red curve). In contrast, (B) shows the increase in the GFP signal (right shift of black curves) corresponding to the SOS induction in the samples exposed to ciprofloxacin when pAPR100 is present. The percentage GFP expressing cells above the negative control was determined by using the inbuilt calculation functions of FCS Express version 7.

## References

1. Frost LS, Leplae R, Summers AO, Toussaint A. Mobile genetic elements: the agents of open source evolution. Nat Rev Microbiol. 2005;3(9):722–32.

2. Norman A, Hansen LH, Sorensen SJ. Conjugative plasmids: vessels of the communal gene pool. Philos Trans R Soc Lond B Biol Sci. 2009;364(1527):2275–89.

3. Wang R, van Dorp L, Shaw LP, Bradley P, Wang Q, Wang X, et al. The global distribution and spread of the mobilized colistin resistance gene mcr-1. Nat Commun. 2018;9(1):1179.

4. Caselli E, D’Accolti M, Soffritti I, Piffanelli M, Mazzacane S. Spread of mcr-1-Driven Colistin Resistance on Hospital Surfaces, Italy. Emerg Infect Dis. 2018;24(9):1752–3.

5. Munoz-Price LS, Poirel L, Bonomo RA, Schwaber MJ, Daikos GL, Cormican M, et al. Clinical epidemiology of the global expansion of Klebsiella pneumoniae carbapenemases. Lancet Infect Dis. 2013;13(9):785–96.

6. Lee CR, Lee JH, Park KS, Kim YB, Jeong BC, Lee SH. Global Dissemination of Carbapenemase-Producing Klebsiella pneumoniae: Epidemiology, Genetic Context, Treatment Options, and Detection Methods. Front Microbiol. 2016;7:895.

7. Pitout JD, Laupland KB. Extended-spectrum beta-lactamase-producing Enterobacteriaceae: an emerging public-health concern. Lancet Infect Dis. 2008;8(3):159–66.

8. Canton R, Coque TM. The CTX-M beta-lactamase pandemic. Curr Opin Microbiol. 2006;9(5):466–75.

9. Canton R, Gonzalez-Alba JM, Galan JC. CTX-M Enzymes: Origin and Diffusion. Front Microbiol. 2012;3:110.

10. Livermore DM, Canton R, Gniadkowski M, Nordmann P, Rossolini GM, Arlet G, et al. CTX-M: changing the face of ESBLs in Europe. J Antimicrob Chemother. 2007;59(2):165–74.

11. Arredondo-Alonso S, Top J, McNally A, Puranen S, Pesonen M, Pensar J, et al. Plasmids Shaped the Recent Emergence of the Major Nosocomial Pathogen Enterococcus faecium. mBio. 2020;11(1).

12. Leon-Sampedro R, DelaFuente J, Diaz-Agero C, Crellen T, Musicha P, Rodriguez-Beltran J, et al. Pervasive transmission of a carbapenem resistance plasmid in the gut microbiota of hospitalized patients. Nat Microbiol. 2021;6(5):606–16.

13. Gumpert H, Kubicek-Sutherland JZ, Porse A, Karami N, Munck C, Linkevicius M, et al. Transfer and Persistence of a Multi-Drug Resistance Plasmid in situ of the Infant Gut Microbiota in the Absence of Antibiotic Treatment. Front Microbiol. 2017;8:1852.

14. Conlan S, Thomas PJ, Deming C, Park M, Lau AF, Dekker JP, et al. Single-molecule sequencing to track plasmid diversity of hospital-associated carbapenemase-producing Enterobacteriaceae. Sci Transl Med. 2014;6(254):254ra126.

15. Hegstad K, Mikalsen T, Coque TM, Werner G, Sundsfjord A. Mobile genetic elements and their contribution to the emergence of antimicrobial resistant Enterococcus faecalis and Enterococcus faecium. Clin Microbiol Infect. 2010;16(6):541–54.

16. San Millan A. Evolution of Plasmid-Mediated Antibiotic Resistance in the Clinical Context. Trends Microbiol. 2018;26(12):978–85.

17. Lopatkin AJ, Meredith HR, Srimani JK, Pfeiffer C, Durrett R, You L. Persistence and reversal of plasmid-mediated antibiotic resistance. Nat Commun. 2017;8(1):1689.

18. Sorensen SJ, Bailey M, Hansen LH, Kroer N, Wuertz S. Studying plasmid horizontal transfer in situ: a critical review. Nat Rev Microbiol. 2005;3(9):700–10.

19. San Millan A, Toll-Riera M, Qi Q, MacLean RC. Interactions between horizontally acquired genes create a fitness cost in Pseudomonas aeruginosa. Nat Commun. 2015;6:6845.

20. Vogwill T, MacLean RC. The genetic basis of the fitness costs of antimicrobial resistance: a meta-analysis approach. Evol Appl. 2015;8(3):284–95.

21. MacLean RC, San Millan A. Microbial Evolution: Towards Resolving the Plasmid Paradox. Curr Biol. 2015;25(17):R764–7.

22. Alonso-Del Valle A, Leon-Sampedro R, Rodriguez-Beltran J, DelaFuente J, Hernandez-Garcia M, Ruiz-Garbajosa P, et al. Variability of plasmid fitness effects contributes to plasmid persistence in bacterial communities. Nat Commun. 2021;12(1):2653.

23. Baker KS, Dallman TJ, Ashton PM, Day M, Hughes G, Crook PD, et al. Intercontinental dissemination of azithromycin-resistant shigellosis through sexual transmission: a cross-sectional study. Lancet Infect Dis. 2015;15(8):913–21.

24. Baker KS, Dallman TJ, Field N, Childs T, Mitchell H, Day M, et al. Horizontal antimicrobial resistance transfer drives epidemics of multiple Shigella species. Nat Commun. 2018;9(1):1462.

25. Gilbart VL, Simms I, Jenkins C, Furegato M, Gobin M, Oliver I, et al. Sex, drugs and smart phone applications: findings from semistructured interviews with men who have sex with men diagnosed with Shigella flexneri 3a in England and Wales. Sex Transm Infect. 2015;91(8):598–602.

26. Ingle DJ, Easton M, Valcanis M, Seemann T, Kwong JC, Stephens N, et al. Co-circulation of Multidrug-resistant Shigella Among Men Who Have Sex With Men in Australia. Clin Infect Dis. 2019;69(9):1535–44.

27. van den Beld MJC, Reubsaet FAG, Pijnacker R, Harpal A, Kuiling S, Heerkens EM, et al. A Multifactorial Approach for Surveillance of Shigella spp. and Entero-Invasive Escherichia coli Is Important for Detecting (Inter)national Clusters. Front Microbiol. 2020;11:564103.

28. Locke RK, Greig DR, Jenkins C, Dallman TJ, Cowley LA. Acquisition and loss of CTX-M plasmids in Shigella species associated with MSM transmission in the UK.Microb Genom. 2021;7(8).

29. Worley JN, Javkar K, Hoffmann M, Hysell K, Garcia-Williams A, Tagg K, et al. Genomic Drivers of Multidrug-Resistant Shigella Affecting Vulnerable Patient Populations in the United States and Abroad. mBio. 2021;12(1).

30. Liao YS, Liu YY, Lo YC, Chiou CS. Azithromycin-Nonsusceptible Shigella flexneri 3a in Men Who Have Sex with Men, Taiwan, 2015-2016. Emerg Infect Dis. 2016;23(2):345–6.

31. Moreno-Mingorance A, Espinal P, Rodriguez V, Goterris L, Fabrega A, Serra-Pladevall J, et al. Circulation of multi-drug-resistant Shigella sonnei and Shigella flexneri among men who have sex with men in Barcelona, Spain, 2015-2019. Int J Antimicrob Agents. 2021;58(3):106378.

32. Hinic V, Seth-Smith H, Stockle M, Goldenberger D, Egli A. First report of sexually transmitted multi-drug resistant Shigella sonnei infections in Switzerland, investigated by whole genome sequencing. Swiss Med Wkly. 2018;148:w14645.

33. Blackwell GA, Hunt M, Malone KM, Lima L, Horesh G, Alako BTF, et al. Exploring bacterial diversity via a curated and searchable snapshot of archived DNA sequences. PLoS Biol. 2021;19(11):e3001421.

34. Petrova V, Chitteni-Pattu S, Drees JC, Inman RB, Cox MM. An SOS inhibitor that binds to free RecA protein: the PsiB protein. Mol Cell. 2009;36(1):121–30.

35. Baharoglu Z, Mazel D. SOS, the formidable strategy of bacteria against aggressions. FEMS Microbiol Rev. 2014;38(6):1126–45.

36. Qin TT, Kang HQ, Ma P, Li PP, Huang LY, Gu B. SOS response and its regulation on the fluoroquinolone resistance. Ann Transl Med. 2015;3(22):358.

37. Williams PCM, Berkley JA. Guidelines for the treatment of dysentery (shigellosis): a systematic review of the evidence. Paediatr Int Child Health. 2018;38(sup1):S50–S65.

38. Murray K, Reddy V, Kornblum JS, Waechter H, Chicaiza LF, Rubinstein I, et al. Increasing Antibiotic Resistance in Shigella spp. from Infected New York City Residents, New York, USA. Emerg Infect Dis. 2017;23(2):332–5.

39. San Millan A, MacLean RC. Fitness Costs of Plasmids: a Limit to Plasmid Transmission. Microbiol Spectr. 2017;5(5).

40. Humphrey B, Thomson NR, Thomas CM, Brooks K, Sanders M, Delsol AA, et al. Fitness of Escherichia coli strains carrying expressed and partially silent IncN and IncP1 plasmids. BMC Microbiol. 2012;12:53.

41. Redondo-Salvo S, Fernandez-Lopez R, Ruiz R, Vielva L, de Toro M, Rocha EPC, et al. Pathways for horizontal gene transfer in bacteria revealed by a global map of their plasmids. Nat Commun. 2020;11(1):3602.

42. Hall JP, Wood AJ, Harrison E, Brockhurst MA. Source-sink plasmid transfer dynamics maintain gene mobility in soil bacterial communities. Proc Natl Acad Sci U S A. 2016;113(29):8260–5.

43. Baltrus DA. Exploring the costs of horizontal gene transfer. Trends Ecol Evol. 2013;28(8):489–95.

44. Hall JPJ, Brockhurst MA, Dytham C, Harrison E. The evolution of plasmid stability: Are infectious transmission and compensatory evolution competing evolutionary trajectories? Plasmid. 2017;91:90–5.

45. Benz F, Huisman JS, Bakkeren E, Herter JA, Stadler T, Ackermann M, et al. Plasmid- and strain-specific factors drive variation in ESBL-plasmid spread in vitro and in vivo. ISME J. 2021;15(3):862–78.

46. Hall JPJ, Wright RCT, Harrison E, Muddiman KJ, Wood AJ, Paterson S, et al. Plasmid fitness costs are caused by specific genetic conflicts enabling resolution by compensatory mutation. PLoS Biol. 2021;19(10):e3001225.

47. Baharoglu Z, Bikard D, Mazel D. Conjugative DNA transfer induces the bacterial SOS response and promotes antibiotic resistance development through integron activation. PLoS Genet. 2010;6(10):e1001165.

48. Golub E, Bailone A, Devoret R. A gene encoding an SOS inhibitor is present in different conjugative plasmids. J Bacteriol. 1988;170(9):4392–4.

49. Smirnova GV, Tyulenev AV, Muzyka NG, Peters MA, Oktyabrsky ON. Ciprofloxacin provokes SOS-dependent changes in respiration and membrane potential and causes alterations in the redox status of Escherichia coli. Res Microbiol. 2017;168(1):64–73.

50. Aziz RK, Bartels D, Best AA, DeJongh M, Disz T, Edwards RA, et al. The RAST Server: rapid annotations using subsystems technology. BMC Genomics. 2008;9:75.

51. Sullivan MJ, Petty NK, Beatson SA. Easyfig: a genome comparison visualizer. Bioinformatics. 2011;27(7):1009–10.

52. Page AJ, Cummins CA, Hunt M, Wong VK, Reuter S, Holden MT, et al. Roary: rapid large-scale prokaryote pan genome analysis. Bioinformatics. 2015;31(22):3691–3.

53. Bortolaia V, Kaas RS, Ruppe E, Roberts MC, Schwarz S, Cattoir V, et al. ResFinder 4.0 for predictions of phenotypes from genotypes. J Antimicrob Chemother. 2020;75(12):3491–500.

54. Sysoeva TA KY, Rodriguez J, Lopatkin AJ, You L. Growth-stage-dependent regulation of conjugation. AIChe Journal. 2020;66(3).

55. Baharoglu Z, Krin E, Mazel D. Connecting environment and genome plasticity in the characterization of transformation-induced SOS regulation and carbon catabolite control of the Vibrio cholerae integron integrase. J Bacteriol. 2012;194(7):1659–67.

56. Sprouffske K, Wagner A. Growthcurver: an R package for obtaining interpretable metrics from microbial growth curves. BMC Bioinformatics. 2016;17:172.

